# Symptom-modulating satRNAs of cucumoviruses affect the orientation and feeding behaviour of *Myzus persicae*

**DOI:** 10.1101/2024.10.02.616216

**Authors:** Barbara Wrzesińska-Krupa, Przemysław Strażyński, Patryk Frąckowiak, Aleksandra Obrępalska-Stęplowska

## Abstract

Plant viruses evolved mechanisms to manipulate host plants to replicate and be efficiently transmitted by insect vectors. In the context of non-persistently transmitted viruses, an important strategy is to change the plant’s metabolism to attract the vectors, and subsequently repel viruliferous insects from the infected plants to facilitate the virus transmission to neighbouring plants. The presence of satellite RNAs (satRNAs), which accompany certain plant RNA virus species and strains, leads to changes in the host plants, such as alterations in the virus-induced infection symptoms, either exacerbating or attenuating them. This study aimed to analyse the influence of satRNAs on the orientation and feeding behaviour of the cucumovirus insect vector – *Myzus persicae*, which might consequently contribute to the virus transmission efficiency. The hypothesis behind this study was that satRNAs of cucumoviruses alter these insect activities toward virus-infected plants, and strongly symptom-deteriorating satRNAs might negatively affect the attractiveness of the infected plants for aphids. Using two cucumoviruses, peanut stunt virus (PSV) and cucumber mosaic virus (CMV), and their satRNAs, which induce divergent infection symptoms, olfactometry and electrical penetration graph (EPG) monitoring analyses were performed. The results showed that satRNAs which presence leads to disease symptom exacerbation might alter the orientation behaviour of aphids by reducing the attractiveness of the plants and discouraging aphids from feeding. This phenomenon may contribute to the better persistence in the environment of satRNAs alleviating disease symptoms compared to the worsening ones, benefiting the virus by not destroying the plant and prolonging the virus’ exposure to insect vectors.

## INTRODUCTION

Plant viruses are intracellular parasites that alter the metabolism of infected plants. They have evolved mechanisms not only to evade plant host antiviral defences but also to ensure their successful replication and movement within the plant and efficient spread between plants (Mauck et al., 2012). An infected plant undergoes such significant metabolic changes that it can become more attractive to insects acting as virus vectors, facilitating the pathogen’s transmission to other plants (Carmo-Sousa et al., 2014, Mauck et al., 2010, Webster, 2012). These changes may concern the physiological and chemical properties of the host plant that condition the attractiveness of the infected plants to vectors by olfactory, visual, and gustatory cues (Mauck et al., 2014, Schröder et al., 2017). Many examples of viruses altering cues expressed by plants in ways that increase transmission, were presented, but there are fewer studies documenting mechanisms, especially for non-persistently transmitted viruses that have only transient interactions with their aphid vectors. Additionally, few studies have considered subviral agents that are transmitted with their helper viruses. These include satellite RNAs (satRNAs), which are capable of modifying host phenotype.

Here, we studied the effects of satRNAs on infections by two non-persistently transmitted viruses in the genus *Cucumovirus*, family *Bromoviridae*: cucumber mosaic virus (CMV) and peanut stunt virus (PSV). CMV infects a wide range of hosts, encompassing more than 1000 plant species, while the host range of PSV mainly involved plants belonging to the Fabaceae and Solanaceae families (Bananej et al., 1998, Jacquemond, 2012). Nevertheless, these virus species are distributed worldwide and cause serious diseases in agriculturally important plants, constituting a significant global problem. Our goal was to determine if satRNAs that modify the host phenotypes induced by their respective helper virus, change vector behaviour in relation to infected hosts. By leading to symptom attenuation, satRNA has a beneficial effect on helper virus transmission, while by inducing symptom aggravation, it restricts helper virus from spreading.

CMV and PSV have positive-sense single-stranded RNA (ss(+)RNA) genomes, consisting of three genomic and two subgenomic strands. RNA1 and RNA2 encode 1a and 2a proteins, respectively, which are the components of the viral replication complex. Additionally, RNA 2 is a source of subgenomic RNA 4A having an open reading frame for the 2b protein, which is involved in RNA silencing suppression and viral movement. Similarly, RNA 3 serves as a translation template for the movement protein (MP, 3a) and coat protein (CP); the latter is synthesized from subgenomic RNA 4 (Ding et al., 1994, Palukaitis et al., 1992). Viral proteins are also engaged in virus-vector-host plant interactions and virus transmissibility. Previous studies documented multiple cases of host and vector manipulation by cucumoviruses, particularly CMV. CP was found to be one of the crucial factors in transmission by the insect vectors. It binds loosely to acrostyle receptors in the insect’s stylet (Webster et al., 2018). The 1a protein of CMV-Fny was shown to trigger *Myzus persicae* (Sulz.) resistance in tobacco, while the 2b protein suppressed the host resistance and altered volatile organic compounds (VOCs) emission in virus-infected plants (Tungadi et al., 2017, Ziebell et al., 2011). On the other hand, the 2a protein causes *M. persicae* feeding deterrence (antixenosis) in *Arabidopsis thaliana*, while the 1a protein counteracts the induction of antibiosis (toxicity) by the 2b protein (Rhee et al., 2020, Westwood et al., 2013).

Some CMV and PSV strains can also be accompanied by satRNA (Simon et al., 2004). Cucumoviral satRNAs do not encode any functional protein and are fully dependent on the helper virus for replication, encapsidation, movement, and transmission (Hu et al., 2009). SatRNAs might influence the symptoms generated by the helper virus (attenuating or exacerbating them), the helper virus accumulation level (usually decreasing) (He et al., 2019, Obrępalska-Stęplowska et al., 2018, Wrzesińska et al., 2018), and pathogenesis progress (Obrępalska-Stęplowska et al., 2013, Xu et al., 2016). The outcome of the satRNA-helper virus-host plant interactions depends on the sequence of satRNA, the strain of the helper virus, the host plant species, and external factors (e.g., temperature) (Wrzesińska Krupa & Obrępalska Stęplowska, 2024). Additionally, recently satRNA was found to alter aphids’ behaviour. CMV Y-sat, by inducing yellow leaf colour in tobacco, was shown to enhance the attractiveness of *M. persicae* toward Y-sat-infected plants through visual cues (Jayasinghe et al., 2021). Moreover, Y-sat caused aphid red colouring and wing formation, subsequently enabling the acceleration of virus transmission to healthy plants despite a lower virus titre in plants infected with CMV Y-sat than in plants infected with the CMV only. However, since satRNAs usually lower CMV titre in the infected plants, they were observed to reduce the efficiency of CMV transmission by *Aphis gossypii* in tomato and melon plants (Betancourt et al., 2011, Escriu et al., 2000). However, we still do not know enough about cucumoviruses other than the well-studied CMV and their interactions with viral vectors. The associations between the severity of disease symptoms due to satRNA co-infection with various virus strains, or co-infection with different satRNAs of the same virus strain, as well as aphids’ orientation and feeding behaviour have never been studied before.

To address this knowledge gap, we studied the behaviour of the green peach aphid, *M. persicae*, against model plant *Nicotiana benthamiana* and a crop plant tomato (*Solanum lycopersicum* L.) infected with CMV and PSV, with or without their satRNAs. We carried out our analyses on two strains of PSV: one naturally devoid of satRNA (strain G, where co-infection of satRNA with this strain leads to symptom exacerbation) and one naturally possessing satRNA (strain P, where co-infection of satRNA with this strain leads to slight symptom attenuation). Additionally, we analysed the CMV Fny strain with non-necrogenic (non-nc-satRNA) and necrogenic satRNAs (nc-satRNAs). We hypothesise that virus effects on olfactory and/or palatability cues are altered by the presence of symptom-modifying satRNAs. Since satRNAs can modify the symptoms of viral infections in plants, leading to their exacerbation or weakening, as well as affecting plants’ metabolism during viral infection, we assume that the presence of satRNA in virus-infected plants modifies plants’ olfactory attractiveness to the insect vector and influences the aphids’ feeding behaviour. The direction of the changes in pathogenesis (lessening or aggravating) caused by satRNA may significantly impact the behaviour of aphids. Since necrogenic satRNAs lead to severe disease symptoms leading to the premature death of a plant, the chances of the virus transmission to neighbouring plants and further multiplication are lowered. In this context, we wondered whether the presence of the satRNA would enhance the attractiveness of plants to insect vectors as long as it does not significantly worsen disease symptoms. We also questioned the species-specificity of the observations, both in relation to the virus and the plant host.

## MATERIALS AND METHODS

### Biological materials

*N. benthamiana* plants used to obtain viral particles from infectious copies were grown under controlled conditions in greenhouse chambers with a 14 h light/10 h dark cycle at 24 °C day/20 °C night. Five-to-six-week-old *N. benthamiana* seedlings were infiltrated with infectious copies of PSV-G or PSV-P, either alone or in combination with satRNA-P (PSV-G + satRNA, PSV-P + satRNA) in *Agrobacterium tumefaciens*. Similarly, plants were infiltrated with infectious copies of CMV-Fny alone or in combinations with non-necrogenic satRNA (CMV + non-nc-satRNA) or necrogenic satRNA (CMV + nc-satRNA). Plants infiltrated with the infiltrating buffer served as negative controls (mock). At 9 days post-infiltration (dpi), the fragments of systemic leaves were harvested for virus and satRNA presence verification. At 14 dpi, systemic leaves from four plants (ca. 2.0 g) were collected and homogenized in 5 ml of 0.05 M phosphate buffer (pH 7.5) using a pestle and mortar. This sap was used for the mechanical inoculation of plants used in subsequent experiments investigating aphid behaviour. Two leaves of 4-week-old *N. benthamiana* (for PSV and CMV) and *S. lycopersicum* cv. Betalux (for CMV but not PSV, as it does not infect tomatoes) plants were dusted with carborundum and inoculated with cucumovirus with or without satRNA combinations (**Figure 1**). Plants treated with the phosphate buffer only (mock) were taken as the negative controls. All plants were grown in a greenhouse chamber with a 14 h light/10 h dark cycle in 21 °C day/18 °C night. For the analysis of aphid orientation preferences (olfactometry), three plants from each treatment (biological replicates) were assessed at 2-3 weeks post-inoculation (wpi), when the infection symptoms began to develop. To monitor aphid feeding behaviour (using an electrical penetration graph, EPG), eight plants from each treatment (biological replicates) were observed both before and after fully developed symptoms. A separate set of plants was prepared for each experiment.

**Figure 1.**
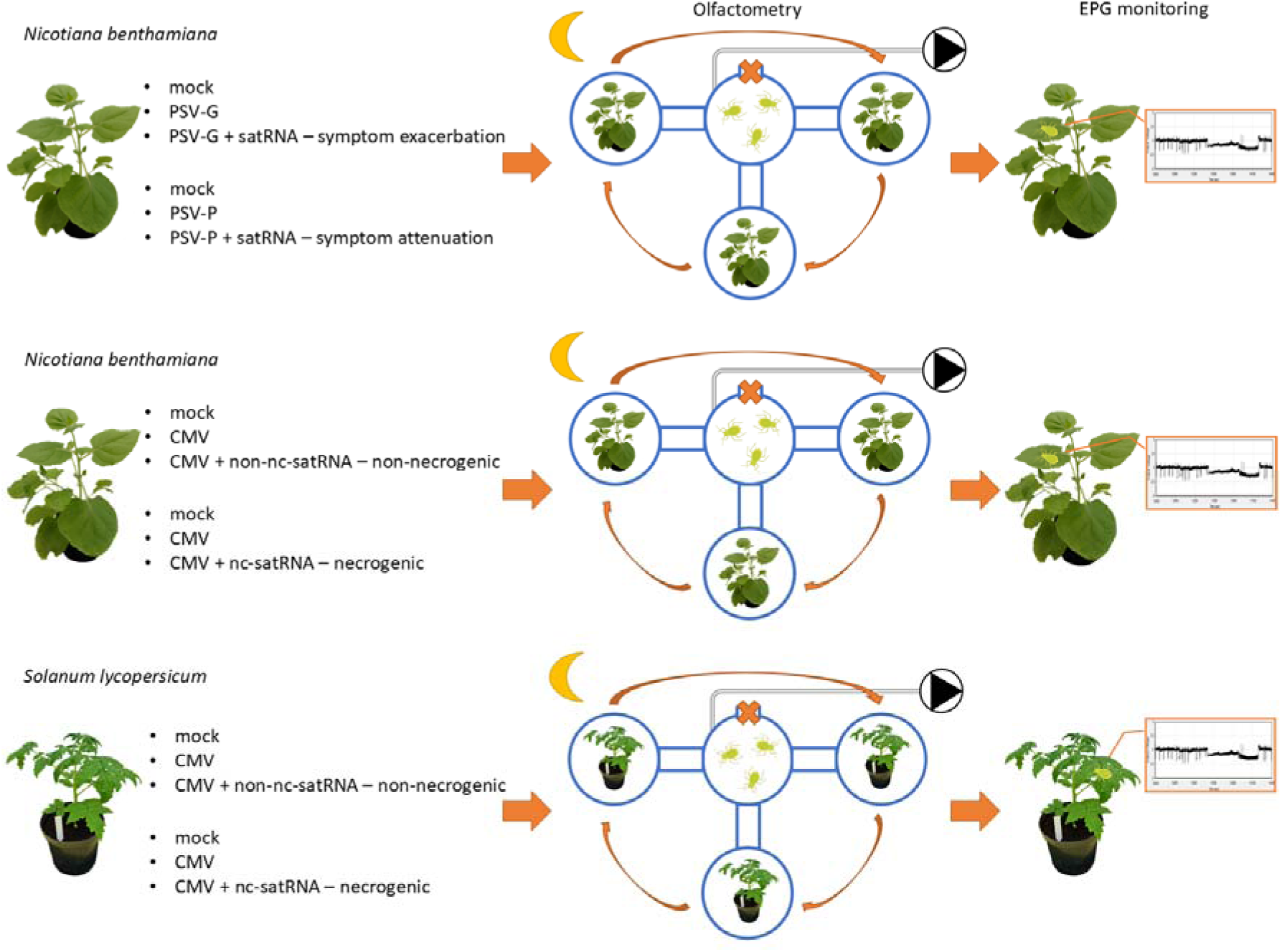
The schematic presentation of the experimental setup to investigate the influence of satellite RNAs (satRNAs) of cucumoviruses on the orientation and gustatory properties of two host plants: *Nicotiana benthamiana* (a model plant), and *Solanum lycopersicum* (a crop plant). Plants infected with the cucumoviruses with or without specific satRNAs were subjected to olfactometer analysis and electrical penetration graph (EPG) monitoring. *N. benthamiana* was infected with peanut stunt virus (PSV) or cucumber mosaic virus (CMV) and *S. lycopersicum* cv. Betalux infected with CMV but not PSV, as it does not infect tomatoes. Plants treated with the phosphate buffer only (mock) were taken as the negative controls. A separate set of plants was prepared for each experiment. PSV-G – PSV strain G, PSV-G + satRNA – PSV strain G with satRNA, PSV-P – PSV strain P, PSV-P + satRNA – PSV strain P with satRNA, CMV + non-nc-satRNAs – CMV and non-necrogenic satRNA, CMV + nc-satRNA – CMV and necrogenic satRNA. An aphid image was obtained from TogoTV

Wingless individuals of *M. persicae* (Sulz.) (peach-potato or green peach aphid) were used in the experiments. The aphid stock colonies were maintained on *N. tabacum* or *Brassica napus* plants placed in an insect cage. The growth conditions were 14 h light/10 h dark cycle in 18-21 °C day/18 °C night.

### Viral infectious clones

These studies involved working with the following viruses: PSV-P, PSV-G, and CMV-Fny. Agroinfectious clones of PSV-P and satRNA-P, and CMV-Fny have been synthesized before (Wrzesińska et al., 2016, Yao et al., 2011). The infectious clones of PSV-G and CMV non-nc-and nc-satRNAs were synthesized as described in Wrzesińska et al. (2016). Previously described infectious clones of PSV-G and cDNA clones of CMV non-nc-and nc-satRNAs cloned under T7 RNA polymerase promoter (Escriu et al., 2000, Obrępalska-Stęplowska et al., 2018) were used as templates for the RNAs amplification through polymerase chain reaction (PCR) with the use forward and reverse primers specified in **Table S1**.

### Virus and satRNA presence verification in the infected plants by reverse transcription polymerase chain reaction (RT-PCR)

To verify the cucumoviruses and satRNAs presence in the agroinfiltrated plants at 9 dpi, 1 cm in diameter leaf fragments were collected for the analysis. Total RNA was extracted using 500 μL of TRI Reagent Solution (Thermo Fisher Scientific, Waltham, MA, USA) followed by precipitation with 2-propanol and washing with 70% ethanol. The air-dried precipitate was suspended in nuclease-free water. The quantity and quality of the RNA were estimated in a DS-11 Series Spectrophotometer (DeNovix, Wilmington, DE, USA). One μg of RNA was reverse transcribed using RevertAid Reverse Transcriptase (Thermo Fisher Scientific) with a random hexamer primer (EURx, Gdańsk, Poland) according to the manufacturer’s instructions. The PCR reaction mixture (10 μL) contained 1× DreamTaq Green PCR Master Mix (Thermo Fisher Scientific), 0.5 μM forward primer, 0.5 μM reverse primer, and 1 μL of cDNA. To verify PSV and satRNA-P presence, PSVCP1 and PSVCP2 primers, and PSVsat1 and PSVsat2 primers were used, respectively (**Table S2)**. Whereas, for CMV as well as non-nc-and nc-satRNAs detection, CMVcpF and CMVcpR primers, and CMVsatF and CMVsatR primers were used, respectively (**Table S2)**.

### Quantification of viral RNAs and satRNAs in the infected plants by quantitative real-time PCR (RT-qPCR)

To assess the cucumoviruses and satRNAs genomic RNA levels in infected plants, leaves from three plants subjected to olfactometer assay were harvested after the analysis (21-23 dpi). Total RNA was extracted as described above, followed by DNA digestion using RNase-free DNase I (Thermo Fisher Scientific). One μg of digested RNA was reverse-transcribed as described above. RT-qPCR was performed in triplicate (technical replicates) for each sample using a LightCycler 480 Instrument (Roche Diagnostics, Basel, Switzerland). The reaction mixtures consisted of iTaq™ Universal SYBR Green Supermix (Bio-Rad, Hercules, California, USA), 0.5 μM forward and reverse primers (**Table S3**), and 1 μL of cDNA. Reactions (qPCR) were performed as follows: 3 min of initial denaturation at 95 °C followed by 40 cycles of 20 s at 95 °C, 20 s annealing (at the temperatures listed in **Table S3**), and 20 s at 72 °C. Dissociation curves were generated during temperature ramping from 65 °C to 95°C.

Absolute quantification was done based on the standard curves created using 5-fold dilutions of cDNA synthesized from RNA isolated from the infected plants. The exact copy number of the analysed corresponding genomic RNAs was calculated using the Avogadro’s number. For the statistical analysis, the normality was checked and Mann-Whitney *U* test was performed. Statistically significant differences were calculated in R (ver. 4.3.2) using the ‘ggstatsplot’ package (ver 0.12.3).

### Analysis of insects’ orientation behaviour using an olfactometer

A 4-arm olfactometer, comprising a central main arena (18 cm in diameter) and four tunnels (15 cm long) leading to side jars (18 cm in diameter), was utilised (**Figure S1**). Each tunnel was terminated with a copper mesh to prevent physical contact between the plant and aphids. To conduct the experiment, plants infected with the virus, virus + satRNA, and mock-inoculated plants were used. One tunnel was covered with parafilm in order to block the aphids from entering. In each side jar, a plant from one treatment was placed and covered with a lid supplemented with a carbon filter to remove any odour from the air intake. Thirty aphids, after 2-3 h of starvation at 22 °C, were placed in the central arena and covered with a lid. An air pump connected to the central jar allowed air intake from the side jars. The olfactometer was covered with a black cloth and kept in a dark room to avoid visual cues. The choice made by the aphids was checked after 20 min. Those that entered the tunnel were considered to have made a choice. Three plants (biological replicates) from each treatment were used. After the test, the positions of the side jars with each biological replicate were swapped to avoid aphid directional cues (three technical replicates per one biological replicate). The number of aphids making a choice in each technical replicate for one plant was then summed. The experiment was conducted twice. The total number of biological replicates was 6. For the statistical analysis, the normality was checked and Kruskal-Wallis test was performed. Statistically significant differences were calculated in R (ver. 4.3.2) using the ‘ggstatsplot’ package (ver. 0.12.3).

### The analysis of the insects’ feeding behaviour using Electrical Penetration Graphs

The feeding of *M. persicae* on *N. benthamiana* and *S. lycopersicum* plants was monitored using the Electrical Penetration Graphs (EPG) technique (Leszczyński & Tjallingii, 1994). The Giga-4 EPG system (Wageningen, Netherlands) was used. A microelectrode (gold wire with a diameter of 12.5 μm and a length of about 2–3 cm) was connected to the wingless aphid female’s body using a drop of silver paint. The aphid was placed on a leaf and monitored for 8 hours. Its feeding behaviour was studied using the EPG Stylet+d computer program. Each recording included a record of the activity of one aphid’s stylets on one plant. Eight plants and eight aphids were used for each treatment (biological replicates) (one aphid per one plant). The EPG Stylet+a registration analysis computer program was used to process the obtained results. The average duration of the following EPG waveforms corresponding to different phases of plant tissue penetration by aphid stylets (Np, ABC, E1, E2) was analyzed. For the statistical analysis, the normality was checked and Mann-Whitney *U* test was performed. Statistically significant differences were calculated in R (ver. 4.3.2) using the ‘ggstatsplot’ package (ver. 0.12.3).

## RESULTS

### *Nicotiana benthamiana* and tomato plants inoculated with infectious copies of the cucumovirus and satRNAs developed characteristic disease symptoms

The infectivity of the generated virus and virus + satRNA infectious clones in the agroinfiltrated *N. benthamiana* plants was confirmed by gel electrophoresis of RT-PCR products resulting from amplification of fragments corresponding to CPs genes and the fragments of satRNAs sequences (**Figure 2**).

**Figure 2.**
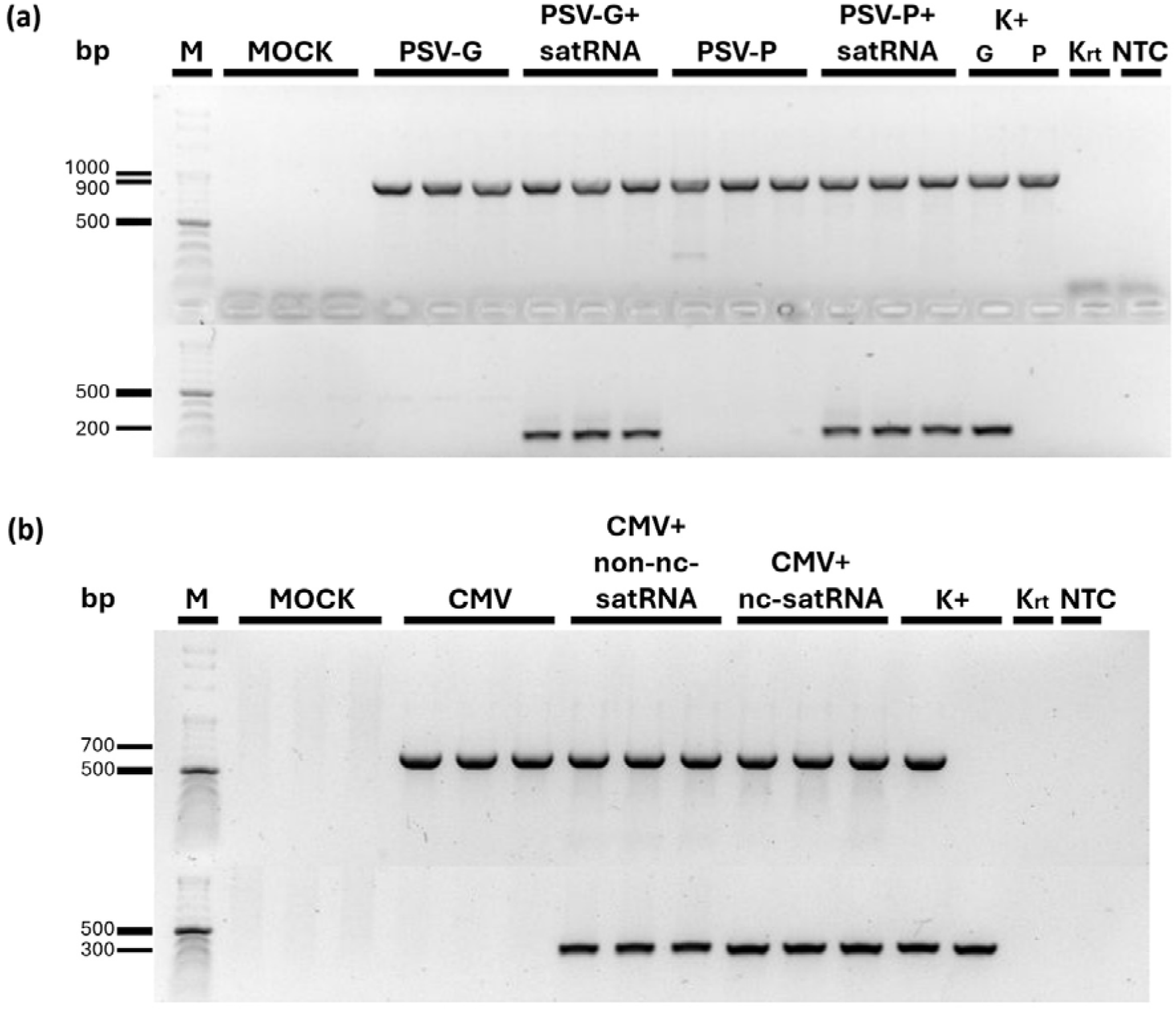
Peanut stunt virus (PSV) (**a**) and cucumber mosaic virus (CMV) (**b**) coat protein (upper row) and satellite RNAs (satRNAs) (lower row) sequences identification in the agroinfiltrated *Nicotiana benthamiana* plants. M – molecular mass marker, K+ – positive controls [plasmids with sequences corresponding to individual RNA strands: in (a) for PSV, PSV-G and PSV-P RNA3 (upper row), and satRNA (lower row); in (b) for CMV, CMV RNA3 (upper row), and non-necrogenic satRNA (non-nc-satRNA) and necrogenic satRNA (nc-satRNA), respectively (lower row)]; Krt – reverse transcription negative control (without RNA template); NTC – no cDNA template control

After the confirmation of the virus and satRNA presence, plants infected with PSV or PSV + satRNA and CMV or CMV + satRNAs were used for passage to *N. benthamiana* (PSV and CMV) and *S. lycopersicum* (CMV). Those plants were used for further olfactometer assay and EPG monitoring. Plants displayed characteristic disease symptoms at 21-23 dpi (**Figure 3**). The infection of *N. benthamiana* plants with PSV-G and PSV-P resulted in reduced plant growth, leaf malformations, and chloroses (**Figure 3a**). SatRNA presence in the PSV-G inoculum led to symptom aggravation, causing a more dwarfed phenotype, more severe systemic leaf distortion, and necroses, while satRNA addition to the PSV-P strain resulted in lessened symptom modifications (**Figure 3a**). CMV infection in *N. benthamiana* resulted in the characteristic symptoms caused by the Fny strain: stunting and leaf malformation, whereas the addition of satRNAs resulted in the emergence of necroses, where nc-satRNA led to a higher number of necrotic spots compared to non-nc-satRNA (**Figure 3b**). On the other hand, *S. lycopersicum* infected with CMV displayed slightly delayed growth, systemic leaf distortion, and shoestring-like apical leaf blades (**Figure 3c**). The presence of non-nc-satRNA led to the attenuation of the infection symptoms to some extent, whereas the presence of nc-satRNA caused the appearance of necroses (**Figure 3c**).

**Figure 3.**
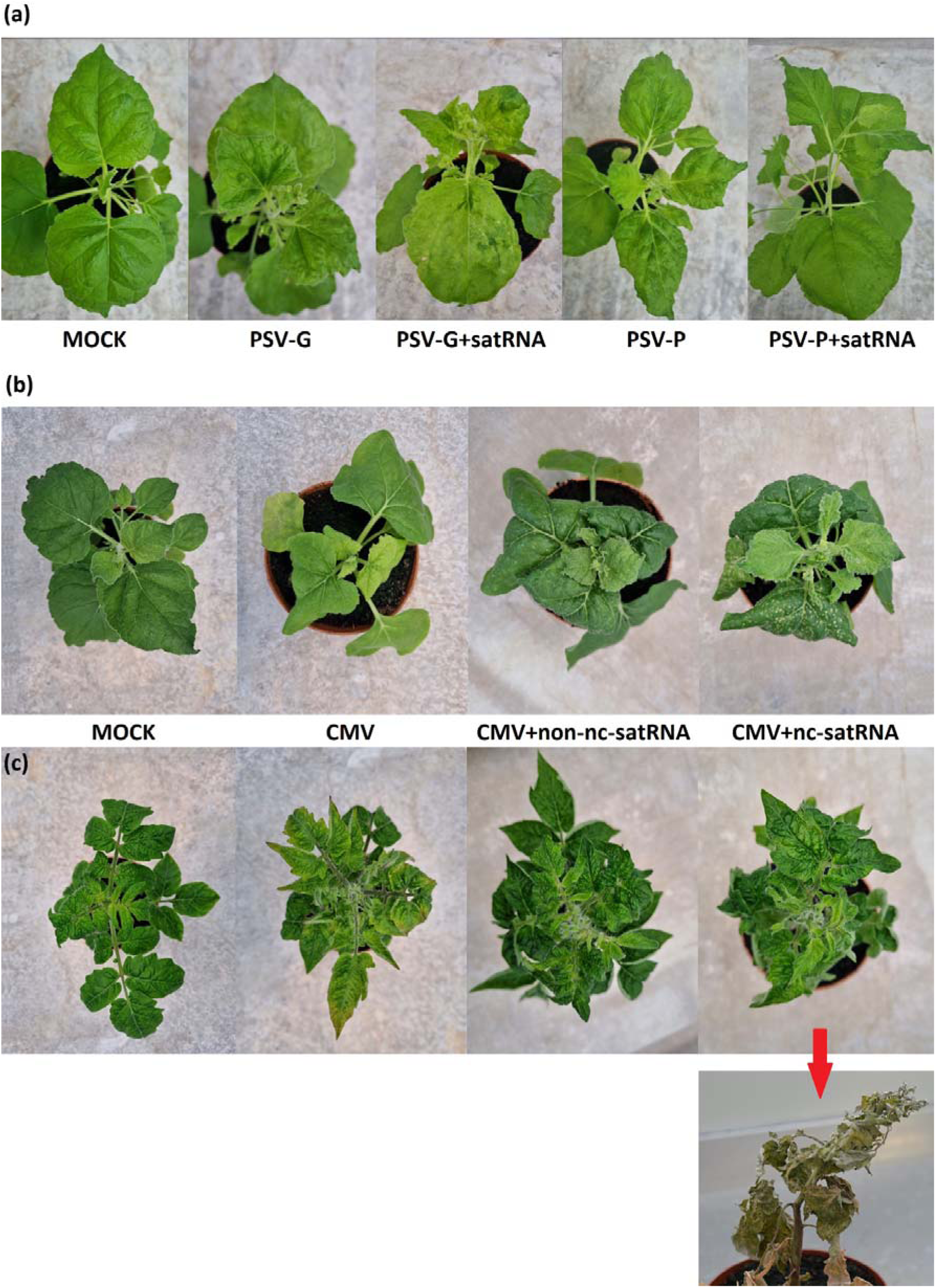
Disease symptoms caused by peanut stunt virus (PSV) or PSV and satellite RNA (PSV + satRNA) on *Nicotiana benthamiana* plants (**a**) and cucumber mosaic virus (CMV) or CMV and non-necrogenic satRNA (CMV + non-nc-satRNA), or CMV and necrogenic satRNA (CMV + nc-satRNA) on *N. benthamiana* (**b**) and *Solanum lycopersicum* (**c**) plants at 21-23 days post-inoculation (dpi) after passage from *N. benthamiana* plants infiltrated with agroinfectious clones of each virus with or without satRNA. An additional photograph of severely developed necrotic symptoms at a later stage of infection (36 dpi) caused by nc-satRNA in CMV inoculum was indicated with red arrow

### SatRNA influences the cucumovirus RNAs’ accumulation levels differently depending on the virus strain, satRNA sequence, and host plant species

Viral genomic strands and satRNAs accumulation analysis was performed on plants subjected to orientation behaviour analysis at 21-23 dpi. The presence of satRNA in the PSV-G inoculum did not significantly alter the accumulation level of PSV genomic strands. However, in plants co-infected with PSV-P with satRNA, the levels of PSV-P RNA1, RNA2, and RNA3 were lower (**Figure 4a**). There was almost no difference in satRNA level between PSV-G + satRNA-and PSV-P + satRNA-infected plants at 3 wpi.

**Figure 4.**
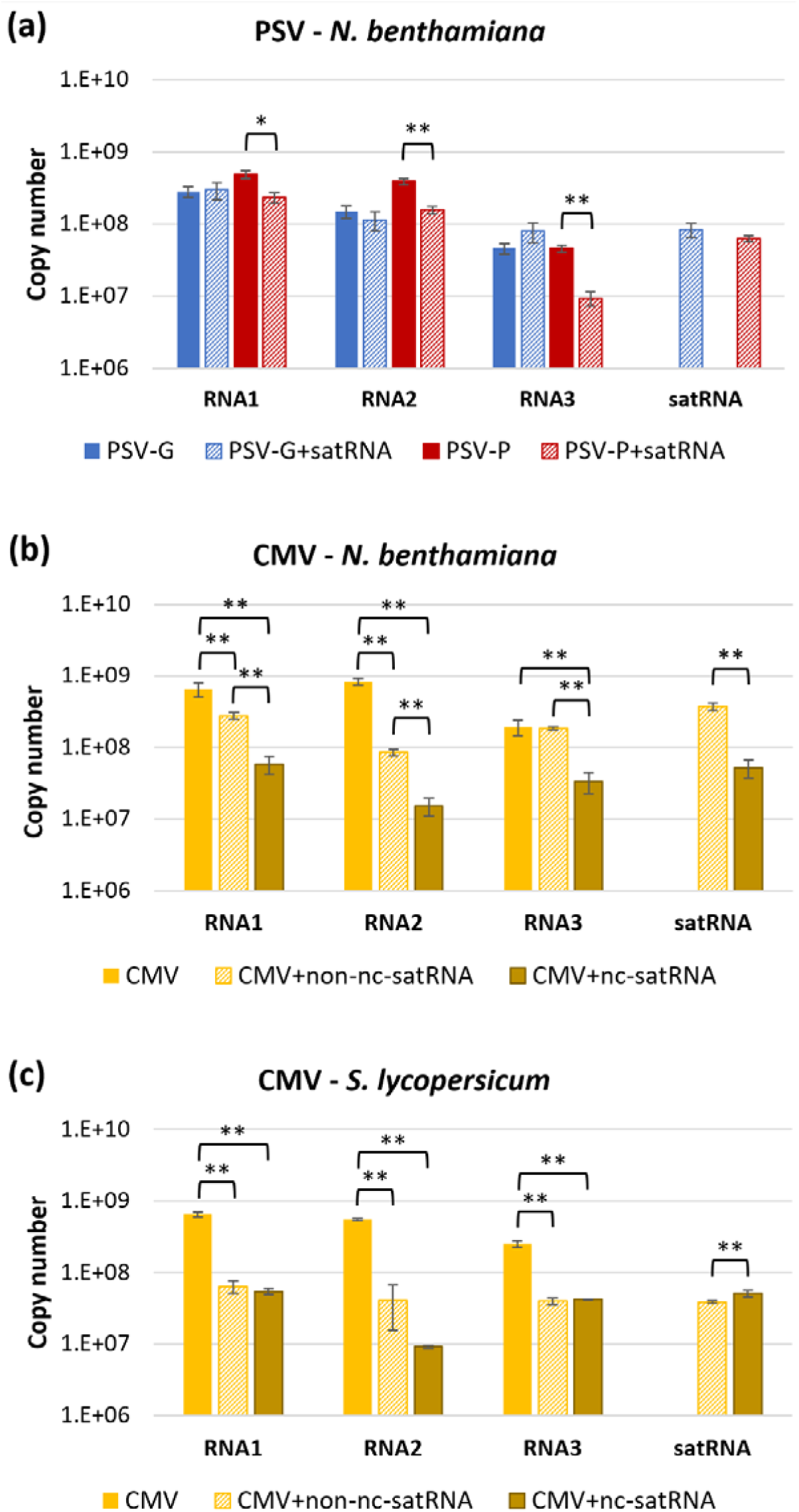
The mean absolute accumulation of cucumoviruses genomic strands [RNA1 (based on 1a gene amplification), RNA2 (based on 2a gene amplification), RNA3 (based on 3a gene amplification)] and their satRNAs. Peanut stunt virus (PSV) accumulation in *Nicotiana benthamiana* plants (**a**) and cucumber mosaic virus (CMV) accumulation in *N. benthamiana* (**b**) and *Solanum lycopersicum* (**c**) plants were assessed at 21-23 days post-inoculation. The reverse transcription-quantitative real-time polymerase chain reaction (RT-qPCR) analysis was performed on three biological with three technical replicates. The copy number was adjusted to the log_10_ scale. Whiskers represent standard error. PSV-G – PSV strain G, PSV-G + satRNA – PSV strain G with satellite RNA (satRNA), PSV-P – PSV strain P, PSV-P + satRNA – PSV strain P with satRNA, CMV + non-nc-satRNAs – CMV and non-necrogenic satRNA, CMV + nc-satRNA – CMV and necrogenic satRNA. * – *p*-value < 0.05, ** – *p*-value < 0.01 (Mann-Whitney *U* test, n = 9). Detailed *p*-values are given in **Table S4** and **Table S5**

Regarding CMV, in both *N. benthamiana* and *S. lycopersicum* infected plants, the presence of satRNAs (non-nc-satRNA and nc-satRNA) significantly caused the reduction in the accumulation of CMV genomic strands (except for RNA3 level in CMV-and CMV + non-nc-satRNA-infected *N. benthamiana*) (**Figure 4b-c**). Nc-satRNA had a more pronounced effect on diminishing the accumulation of CMV genomic RNAs compared to non-nc-satRNA in *N. benthamiana*. (**Figure 4b**). In *S. lycopersicum*, there were no significant differences in CMV genomic strands’ accumulation between CMV + non-nc-satRNA-and CMV + nc-satRNA-infected plants (except for lower accumulation of RNA2 without statistical significance). Additionally, in this plant species, nc-satRNA accumulated 1.3-fold higher than non-nc-satRNA (**Figure 4c**).

### Co-infection of virus-infected plants with satRNA alters the orientation behaviour of aphids

To examine whether satRNA affects plant attractiveness to aphids, a 3-arm olfactometer was utilised to allow aphids to select between non-infected, cucumovirus-infected, and cucumovirus + satRNA-infected plants. Only ∼16% and ∼18% of *M. persicae* have made any choice between non-infected, PSV-, or PSV + satRNA-infected *N. benthamiana* for PSV-G and PSV-P strains, respectively. The analysis of the selection by *M. persicae* between mock-, PSV-G-, and PSV-G + satRNA-inoculated *N. benthamiana* plants revealed that aphids were mostly attracted to plants infected with the virus, followed by plants infected with the virus + satRNA, and least attracted to the non-infected plants (**Figure 5a**). Additionally, a statistically significant preference was detected between PSV-G-and mock-infected *N. benthamiana*, showing that plants infected with the virus only (where symptoms are less severe compared to plants inoculated with PSV-G + satRNA) were the most attractive for aphids. On the other hand, the analysis of the selection by *M. persicae* between mock-, PSV-P-, and PSV-P + satRNA-infected *N. benthamiana* plants revealed that aphids were most attracted to plants infected with the virus + satRNA, followed by mock-and PSV-P-inoculated plants, which drew a similar mean number of aphids (**Figure 5a**). A statistically significant difference was detected between plants inoculated with PSV-P and PSV-P + satRNA, indicating a higher attraction of aphids toward plants with attenuated symptoms.

**Figure 5.**
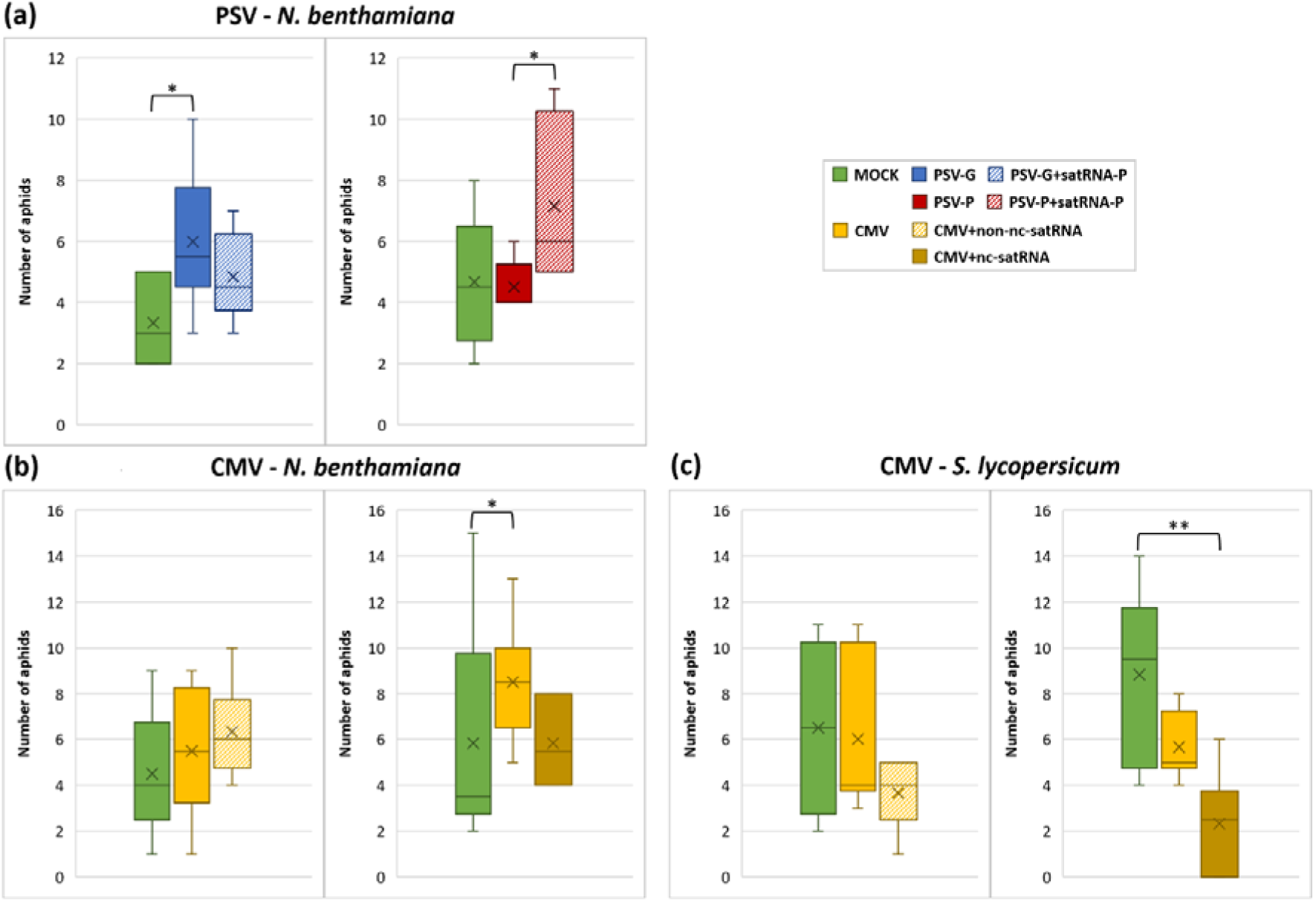
Box plots showing the number of the *Myzus persicae* individuals attracted to the non-infected (mock-inoculated), virus-infected, or virus + satellite RNA (satRNA)-infected plants in a 3-arm olfactometer. The summed number of aphids from three technical replicates with the use of one biological replicate was shown. There were 6 biological replicates. The analysis was done with the use of *Nicotiana benthamiana* plants infected with peanut stunt virus (PSV) or PSV with satRNA (PSV + satRNA) (**a**), *N. benthamiana* plants infected with CMV or CMV + satRNAs (**b**) and *Solanum lycopersicum* plants infected with CMV or CMV + satRNAs (**c)**. Significant differences were calculated according to Kruskal-Wallis test (n = 6). The pairwise comparisons were done with the Dunn test: * – *p*-value < 0.05, ** – *p*-value < 0.01, × and — – mean and median number of aphids out of 90 individuals attracted to the non-infected (mock-inoculated), virus-infected, or virus + satRNA-infected plants. PSV-G – PSV strain G, PSV-G + satRNA – PSV strain G with satellite RNA (satRNA), PSV-P – PSV strain P, PSV-P + satRNA – PSV strain P with satRNA, CMV + non-nc-satRNAs – CMV and non-necrogenic satRNA, CMV + nc-satRNA – CMV and necrogenic satRNA

In the case of the analysis of non-infected, CMV-infected, or CMV + non-nc-satRNA-infected *N. benthamiana*, ∼15% of aphids were attracted to any of the analysed plants. While, in the pathosystem: non-infected, CMV-infected, or CMV + nc-satRNA-infected, this percentage was ∼22%. Among the aphids that made a choice, a greater mean number of individuals preferred CMV-infected plants over mock-inoculated ones (**Figure 5b**). Statistically significant differences (*p*-value < 0.05) were detected when aphids were about to choose between mock-, CMV, or CMV + nc-satRNA-infected *N. benthamiana*. On average, more aphids were attracted to CMV + non-nc-satRNA-infected plants, while fewer individuals chose CMV + nc-satRNA-inoculated *N. benthamiana* compared to those infected with CMV. However, these differences in aphids’ choice between CMV alone and CMV + satRNAs (both non-nc-satRNA and nc-satRNA) were not statistically significant.

Regarding the analysis of *S. lycopersicum* plants – non-infected, CMV-infected, or CMV + non-nc-satRNA-infected – ∼18% of aphids were attracted to any of the analysed plants. In the pathosystems involving non-infected, CMV-infected, or CMV + nc-satRNA-infected plants, this percentage was ∼19%. Among the aphids that made a choice, a higher mean number of *M. persicae* individuals moved toward non-infected *S. lycopersicum* compared to CMV-inoculated plants (**Figure 5c**). The presence of CMV + non-nc-satRNA or CMV + nc-satRNA attracted the fewest aphids, although statistically significant differences were only observed for the comparison of mock-infected *vs* CMV + nc-satRNA-infected plants (*p*-value < 0.01). The comparison between CMV and CMV + non-nc-satRNA or CMV + nc-satRNA did not result in significant differences.

The presence of symptom-alleviating satRNAs in viral inocula enhanced the attraction of aphids towards co-infected plants, while the symptom-exacerbating satRNAs, such as satRNA in PSV-G and nc-satRNA in CMV, had a discouraging effect on aphids’ preferences. Although the differences between a few comparisons were not statistically significant, presumably due to sample size or due to the low percentage of aphids making any choice, this trend in the change of aphid orientation behaviour is noticeable.

### *Myzus persicae* displays variable feeding activity depending on cucumovirus species and host plant species, which may be modulated by satRNA presence

The analysis of EPG waveforms aimed to verify whether satRNA influences the behaviour of *M. persicae* during feeding on the cucumovirus-infected plants. The analysis was focused on the total time of 4 phases of aphid feeding activity: non-penetration (Np), penetration of peripheral tissues (mesophyll) (ABC), accessing phloem elements and secretion of saliva (E1), and ingestion of phloem sap (E2). Measurements were taken at two time points: the early phase of infection, before symptom development, and the later phase, when symptoms appeared on the infected plants. Although the aphids were monitored for 8 hours, their feeding activity lasted for approximately 2 hours. The time before the aphids started feeding and after they finish feeding was not taken into account.

When considering all analysed conditions together (virus strain, type of satRNA, and host plant species), the Np waveform was consistently shortest when aphids fed on the healthy plants. It became longer during feeding on virus-and virus + satRNA-infected plants. In some cases, this phase was even more elongated during feeding on virus + satRNA-infected plants. However, concerned the subsequent phases of feeding showed varied observations. Before disease symptom development, aphids feeding on both PSV and PSV + satRNA-infected plants had reduced feeding activity (no penetration). However, those feeding on PSV + satRNA-inoculated *N. benthamiana* were the least active (with statistically significant differences detected between insects placed on PSV-G + satRNA-and mock-inoculated plants) (**Table 1**). The same trend persisted after symptom development, with significant variation in the total duration of the Np waveform observed in almost all comparisons (**Table 1**). The increased total time of the Np waveform correlated with the shortening of the remaining waveforms.

**Table 1.**
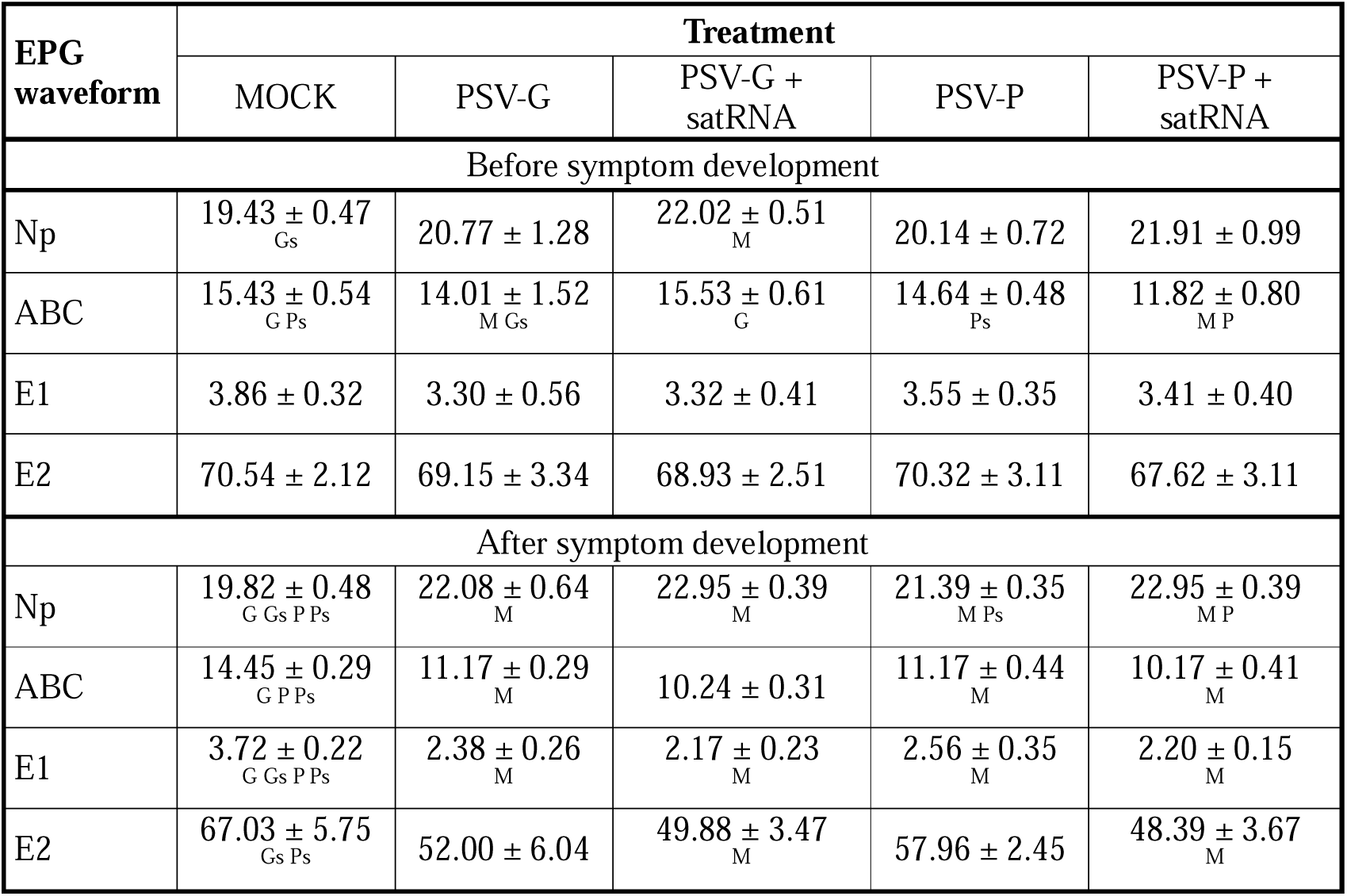
Total duration (minutes, mean ± SEM) of electrical penetration graph (EPG) waveforms during *Myzus persicae* feeding on *Nicotiana benthamiana* plants infected with PSV strains (strain G or strain P) with or without satRNA before and after symptom development. Eight plants and eight aphids were used for each treatment (biological replicates) (one aphid per one plant) (n = 8). Significant differences (*p*-value < 0.05) according to Mann-Whitney *U* test between ‘Treatment’ and: ^M^ – MOCK, ^G^ – PSV-G, ^Gs^ – PSV-G + satRNA, ^P^ – PSV-P, ^Ps^ – PSV-P + satRNA. Detailed *p*-values are given in **Table S6**. Np -non-penetration, ABC – penetration of peripheral tissues (mesophyll), E1 – accessing phloem elements and secretion of saliva, E2 – ingestion of phloem sap

| EPG waveform | Treatment |  |  |  |  |
| --- | --- | --- | --- | --- | --- |
|  | MOCK | PSV-G | PSV-G + satRNA | PSV-P | PSV-P + satRNA |
| Before symptom development |  |  |  |  |  |
| Np | 19.43 ± 0.47<sub>Gs</sub> | 20.77 ± 1.28 | 22.02 ± 0.51<sub>M</sub> | 20.14 ± 0.72 | 21.91 ± 0.99 |
| ABC | 15.43 ± 0.54<sub>G Ps</sub> | 14.01 ± 1.52<sub>M Gs</sub> | 15.53 ± 0.61<sub>G</sub> | 14.64 ± 0.48<sub>Ps</sub> | 11.82 ± 0.80<sub>M P</sub> |
| E1 | 3.86 ± 0.32 | 3.30 ± 0.56 | 3.32 ± 0.41 | 3.55 ± 0.35 | 3.41 ± 0.40 |
| E2 | 70.54 ± 2.12 | 69.15 ± 3.34 | 68.93 ± 2.51 | 70.32 ± 3.11 | 67.62 ± 3.11 |
| After symptom development |  |  |  |  |  |
| Np | 19.82 ± 0.48<sub>G Gs P Ps</sub> | 22.08 ± 0.64<sub>M</sub> | 22.95 ± 0.39<sub>M</sub> | 21.39 ± 0.35<sub>M Ps</sub> | 22.95 ± 0.39<sub>M P</sub> |
| ABC | 14.45 ± 0.29<sub>G P Ps</sub> | 11.17 ± 0.29<sub>M</sub> | 10.24 ± 0.31 | 11.17 ± 0.44<sub>M</sub> | 10.17 ± 0.41<sub>M</sub> |
| E1 | 3.72 ± 0.22<sub>G Gs P Ps</sub> | 2.38 ± 0.26<sub>M</sub> | 2.17 ± 0.23<sub>M</sub> | 2.56 ± 0.35<sub>M</sub> | 2.20 ± 0.15<sub>M</sub> |
| E2 | 67.03 ± 5.75<sub>Gs Ps</sub> | 52.00 ± 6.04 | 49.88 ± 3.47<sub>M</sub> | 57.96 ± 2.45 | 48.39 ± 3.67<sub>M</sub> |

The total duration of ABC waveform before symptom development was significantly shorter for PSV-infected plants that exhibited alleviated symptoms, specifically those inoculated with PSV-G and PSV-P + satRNA (**Table 1**). However, after symptom development aphids devoted less time penetrating mesophyll tissues on plants infected with PSV and even less time on PSV + satRNA-inoculated ones (regardless of virus strain) (**Table 1**). A similar trend was observed for the subsequent E1 and E2 waveforms. However, this clear tendency was not observed before symptom development.

The data relating to the feeding behaviour of aphids on CMV-or CMV + satRNA-inoculated *N. benthamiana* plants are different. Aphids feeding on plants infected with CMV or CMV + satRNAs had longer total Np waveform duration, where the presence of nc-satRNA increases the duration of this waveform the most before and after symptom development (**Table 2**). The total time of ABC waveform is slightly shorter on CMV-infected plants, but longer in the presence of both satRNAs when compared to mock-inoculated *N. benthamiana* before symptom development. However, after symptom development, the time of ABC phase duration was shorter as the infection symptoms were more aggravated (with the shortest for CMV + nc-satRNA) (**Table 2**). Likewise, the total times of E1 and E2 waveforms were similar accordingly to symptom development. However, the only statistically significant differences between the total duration of EPG waveforms concerned the comparison between the total time spent on Np and E1 waveforms during feeding on CMV + nc-satRNA-infected and mock-inoculated plants after symptom development (**Table 2**). This suggests reduced feeding activity of aphids caused by the presence of nc-satRNA.

**Table 2.** Total duration (minutes, mean ± SEM) of electrical penetration graph (EPG) waveforms during *Myzus persicae* feeding on *Nicotiana benthamiana* plants infected with cucumber mosaic virus (CMV) with or without non-necrogenic satRNA (non-nc-satRNA) or necrogenic satRNA (nc-satRNA) before and after symptom development. Eight plants and eight aphids were used for each treatment (biological replicates) (one aphid per one plant) (n = 8). Significant differences (*p*-value < 0.05) according to Mann-Whitney *U* test between ‘Treatment’ and: ^M^ – MOCK, ^C^ – CMV, ^Cnonnc^ – CMV + non-nc-satRNA, ^Cnc^ – CMV + nc-satRNA. Detailed *p*-values are given in **Table S7**. Np -non-penetration, ABC – penetration of peripheral tissues (mesophyll), E1 – accessing phloem elements and secretion of saliva, E2 – ingestion of phloem sap

| EPG waveform | Treatment |  |  |  |
| --- | --- | --- | --- | --- |
|  | MOCK | CMV | CMV + non-nc-satRNA | CMV + nc-satRNA |
| Before symptom development |  |  |  |  |
| Np | 20.59 ± 1.80 | 22.03 ± 2.69 | 23.66 ± 1.95 | 24.60 ± 2.85 |
| ABC | 19.16 ± 1.67 | 18.57 ± 1.05 | 20.12 ± 1.64 | 21.89 ± 1.81 |
| E1 | 4.09 ± 0.49 | 4.31 ± 0.83 | 3.91 ± 0.63 | 3.20 ± 0.68 |
| E2 | 41.89 ± 4.09 | 41.00 ± 4.12 | 37.23 ± 4.29 | 34.09 ± 4.37 |
| After symptom development |  |  |  |  |
| Np | 21.84 ± 0.72<br><sub>Cnc</sub> | 25.04 ± 1.81 | 26.37 ± 1.85 | 27.54 ± 1.70<br><sub>M</sub> |
| ABC | 20.25 ± 0.71 | 19.09 ± 1.52 | 18.81 ± 0.91 | 17.11 ± 1.95 |
| E1 | 3.01 ± 0.43<br><sub>Cnc</sub> | 2.66 ± 0.51 | 1.92 ± 0.31 | 1.60 ± 0.26<br><sub>M</sub> |
| E2 | 40.16 ± 3.18 | 39.38 ± 2.41 | 36.49 ± 2.79 | 32.25 ± 3.91 |

Regarding the feeding behaviour of *M. persicae* on *S. lycopersicum* inoculated with CMV or CMV + satRNAs, the aphids were the least active on CMV + non-nc-satRNA before symptom development. However, the total duration of the Np waveform was longer as the symptoms of infection worsened after symptom development (with the longest duration observed for CMV + nc-satRNA) (**Table 3**). The total time of the ABC waveform was slightly shorter for aphids feeding on CMV-infected plants but longer in the presence of both satRNAs when compared to insects feeding on mock-inoculated *S. lycopersicum* before symptom development. After symptom development, infection with CMV and CMV + satRNAs led to the elongation of the total time of the ABC waveform, where the time of penetration of mesophyll tissues was the longest for aphids feeding on CMV + non-nc-satRNA-inoculated plants (**Table 3**). Additionally, the highest reduction in the total time of phloem waveforms (E1 and E2) was also noted for *M. persicae* feeding on CMV + non-nc-satRNA-infected *S. lycopersicum* before and after symptom development. Furthermore, the total time of the E2 waveform was also the shortest for aphids feeding on CMV + non-nc-satRNA-infected plants before symptom development. However, after symptoms emerged, insects feeding on CMV + nc-satRNA-infected plants collected phloem sap for the shortest time, although the difference was also noticeable for CMV + non-nc-satRNA (**Table 3**). This significant reduction in total phloem waveforms (E1 and E2) times during the 8-hour monitoring of aphids feeding on both CMV + non-nc-satRNA-and CMV + nc-satRNA-infected plants resulted from no activity occurring in these phases by several individuals (**Table 4**). All aphids were engaged in the E1 waveform on CMV-infected plants, while one did not exhibit the E2 waveform. On the other hand, the co-infection of CMV with satRNAs reduced the engagement of *M. persicae* in the phloem phase, where a higher number of individuals did not proceed to the collection of phloem sap from CMV + nc-satRNA-infected plants (**Table 4**). However, there were no statistically significant differences in the total duration of each EPG waveform during *M. persicae* feeding on *S. lycopersicum* infected with CMV or CMV + satRNAs (**Table 3**).

**Table 3.** Total duration (minutes, mean ± SEM) of electrical penetration graph (EPG) waveforms during *Myzus persicae* feeding on *Solanum lycopersicum* plants infected with cucumber mosaic virus (CMV) with or without non-necrogenic satRNA (non-nc-satRNA) or necrogenic satRNA (nc-satRNA) before and after symptom development. Eight plants and eight aphids were used for each treatment (biological replicates) (one aphid per one plant) (n = 8). Significant differences (*p*-value < 0.05) according to Mann-Whitney *U* test between ‘Treatment’ and: ^M^ – MOCK, ^C^ – CMV, ^Cnonnc^ – CMV + non-nc-satRNA, ^Cnc^ – CMV + nc-satRNA. Detailed *p*-values are given in **Table S8**. Np -non-penetration, ABC – penetration of peripheral tissues (mesophyll), E1 – accessing phloem elements and secretion of saliva, E2 – ingestion of phloem sap

| EPG waveform | Treatment |  |  |  |
| --- | --- | --- | --- | --- |
|  | MOCK | CMV | CMV + non-nc-satRNA | CMV + nc-satRNA |
| Before symptom development |  |  |  |  |
| Np | 33.22 ± 1.77 | 34.05 ± 2.01 | 36.11 ± 1.89 | 34.61 ± 1.96 |
| ABC | 24.02 ± 1.12 | 23.29 ± 1.13 | 25.02 ± 0.96 | 24.98 ± 1.14 |
| E1 | 2.66 ± 0.30 | 2.53 ± 0.35 | 2.16 ± 0.32 | 2.23 ± 0.32 |
| E2 | 29.83 ± 1.06 | 28.19 ± 1.52 | 26.58 ± 1.81 | 27.47 ± 0.98 |
| After symptom development |  |  |  |  |
| Np | 32.61 ± 2.54 | 35.19 ± 1.67 | 37.36 ± 2.69 | 39.70 ± 4.3 |
| ABC | 29.88 ± 1.94 | 32.39 ± 2.67 | 34.97 ± 3.35 | 33.58 ± 2.27 |
| E1 | 1.21 ± 0.25 | 1.35 ± 0.37 | 0.91 ± 0.38 | 1.14 ± 0.39 |
| E2 | 20.01 ± 3.94 | 23.49 ± 4.58 | 12.87 ± 5.19 | 10.13 ± 4.83 |

**Table 4.** Number of *Myzus persicae* individuals engaged and not engaged in the phloem phase (PP) (E1 and E2 waveforms) during feeding on *Solanum lycopersicum* plants infected with cucumber mosaic virus (CMV), or CMV in co-infection with non-necrogenic satRNA (CMV + non-nc-satRNA), or CMV in co-infection with necrogenic satRNA (CMV + nc-satRNA) after symptom development

| Treatment | E1 |  | E2 |  |
| --- | --- | --- | --- | --- |
|  | engaged in PP | not engaged in PP | engaged in PP | not engaged in PP |
| CMV | 8 | 0 | 7 | 1 |
| CMV + non-nc-satRNA | 5 | 3 | 4 | 4 |
| CMV + nc-satRNA | 6 | 2 | 3 | 5 |

## DISCUSSION

The evolutionary success of crop viruses is largely due to their efficient transmission by vectors. Previous research indicated that virus-infected plants can emit signals that attract insects, which then settle on plants, acquire viral particles, and spread them to subsequent hosts (Mauck et al., 2010). Interestingly, while these signals are generally attractive to insects, in the case of aphids that transmit viruses in a non-persistent manner, the host plant palatability usually deteriorates (Donnelly et al., 2019, Mauck et al., 2010, 2014). After tasting the infected plant (thus acquiring the virus *via* the stylet in a very short time), the aphid moves to probe other plants, thereby transferring the virus (Carmo-Sousa et al., 2014, Donnelly et al., 2019, Westwood et al., 2013). The presence of satRNAs in co-infection with their helper viruses is a potential contributor to changes in the efficiency of virus transmission through insects. For instance, recent studies have shown that the presence of Y-sat during CMV infection, despite lower virus levels and change in symptom expression on CMV-infected plants, does not affect olfactory attractiveness for aphids (Jayasinghe et al., 2021). However, more aphids that fed on CMV + Y-satRNA-co-infected plants displayed winged morphs than those that fed on CMV-only infected plants. The presence of such satRNA therefore enables the spread of the virus in the environment over longer distances.

In this study, we wondered whether satRNAs may have a discouraging effect on insects’ behaviour (orientation and feeding) when co-infected with some virus strains they cause a strong deterioration of infection symptoms. In consequence, this situation is unfavourable for the host but may also be disadvantageous for virus transmission. Therefore, they may contribute to shortening the exposure of the virus to insects. Perhaps, as a consequence, there may be a reduction in the population of such satRNAs in the environment. Specifically, we focused on two cucumoviruses, PSV and CMV, with which satRNAs exacerbate symptoms or not. We investigated whether the appearance of the necrogenicity or significant symptom deterioration caused by satRNA affects the orientation and feeding behaviour of insects on plants co-infected with cucumovirus and satRNA.

In the case of the interactions between *M. persicae* and *N. benthamiana* infected with two strains of PSV (with or without satRNA), the presence of satRNA in the PSV-G inoculum led to symptom exacerbation and the emergence of necroses. SatRNA presence caused slight changes in the accumulation of PSV-G genomic strands. These changes were not statistically significant, however, the direction of these alterations was in accordance with results reported by Obrępalska-Stęplowska et al. (2018). On the other hand, satRNA co-infection with PSV-P in *N. benthamiana*, where symptom development is slightly attenuated, the accumulation level of helper virus genomic strands was diminished concordantly with data described by Wrzesińska et al. (2018). CMV-Fny resulted in severe symptoms of infection in both *N. benthamiana* and *S. lycopersicum* plants. However, despite a lower accumulation level of viral genomic strains resulting from the presence of non-nc-satRNA and nc-satRNA, the symptoms remained exacerbated in *N. benthamiana*, with more profound necroses during infection with CMV + nc-satRNA. While, in *S. lycopersicum*, the symptoms caused by CMV + non-nc-satRNA were slightly attenuated but infection with CMV + nc-satRNA led to the emergence of severe necroses. Considerable reduction in CMV RNA accumulation caused by non-nc-satRNAs and nc-satRNAs was also indicated before (Betancourt et al., 2011, Escriu et al., 2000).

Our results showed that depending on the host plant, virus species, and satRNA, the behaviour of aphids differs, which is in line with previous studies. The analysis of orientation behaviour revealed that symptom-deteriorating satRNAs of both cucumoviruses reduced the attractiveness of the host plants in relation to plants infected only with the virus (*N. benthamiana* in case of PSV-G + satRNA and CMV + nc-satRNA, and *S. lycopersicum* in case of CMV + nc-satRNA) for *M. persicae* individuals, which confirms our hypothesis. However, a few results were not statistically significant. Nevertheless, the addition of satRNA to viral inoculum that causes necroses can lead to a faster death of the plant and shorten the possibilities of the attraction of aphids to those plants and subsequent virus transmission to the healthy plants.

Aphids significantly preferred *N. benthamiana* plants infected with PSV-G (strain naturally devoid of satRNA) and CMV over mock-inoculated plants. There was no evident difference between the preferences of aphids for CMV-or mock-inoculated *S. lycopersicum*. On the other hand, in the other olfactometry assay, *M. persicae* preferred Fny-CMV-infected *S. lycopersicum* cv. Moneymaker over mock-inoculated (Arinaitwe et al., 2022). Differences between these studies and ours may also result from the use of different *S. lycopersicum* varieties, which may be different in terms of metabolism and susceptibility to viral infections.

In the case of symptom-alleviating satRNAs, the presence of satRNA in PSV-P inoculum, naturally occurring in the environment, despite mitigating symptoms, attracted a higher number of aphids. Similarly, more aphids preferred CMV + non-nc-satRNA-infected *N. benthamiana* plants, but a reduction in aphid attraction was observed for CMV + non-nc-satRNA-inoculated *S. lycopersicum* plants. However, no difference in the aphid’s orientation behaviour caused by satRNA presence was reported in the case of tobacco plants infected with CMV + Y-satRNA (Jayasinghe et al., 2021). Based on our results, the presence of symptom-attenuating satRNAs may be beneficial for virus survival in the environment, as the host plant endures the infection despite efficient multiplication of helper virus, thereby ensuring exposure of the virus to aphid acquisition and its transmission to subsequent plants.

The acquisition of the non-persistently transmitted viruses by stylet penetrations of epidermal cells of the infected leaves requires a very short time. This strategy enables efficient uptake of viral particles (Donnelly et al., 2019). Based on the previous observations, infection with non-persistently transmitted viruses can increase the attractiveness of the host plants for aphids and encourage them to probe the infected leaves (corresponding to the ABC waveform) (Mauck et al., 2012, Mauck et al., 2010). However, the palatability of the infected plants deteriorates and the feeding time of aphids shortens (corresponding to the E1and E2 waveforms), causing aphids to leave the plants, which in consequence promotes faster dispersion of the virus in the environment (Mauck et al., 2012, Mauck et al., 2010, 2014). In most cases, we observed a trend of elongation of the non-probing phase and shortening of the phloem phase caused by the infection process. Furthermore, this tendency was more pronounced when plants were infected with virus + satRNA. SatRNA in co-infection with both PSV-G and PSV-P reduced the total time of ABC waveform after symptoms development on *N. benthamiana* plants. Moreover, there was also a significant reduction in the duration of the phloem phase (E1 and E2 waveform) when aphids ingested sap from PSV and satRNA-infected *N. benthamiana* for both G and P strains. It can be concluded that satRNA co-infection with PSV shortens the duration of this phase regardless of symptoms severity (exacerbation in PSV-G + satRNA-and alleviation in PSV-P + satRNA-infected plants), which increases the chances of aphids escaping from these plants and transmitting the virus to healthy plants. These data overlap with the orientation behaviour analysis suggesting “attract and deter” manipulation by PSV (and satRNA), in which aphids are attracted to the infected plants by olfactory cues and repelled by gustatory cues to efficiently infect the adjacent plants (Donnelly et al., 2019, Mauck et al., 2010).

The time that aphids spent on probing and phloem phase varied between *N. benthamiana* and *S. lycopersicum* plants. A ratio between the means of total times of the E2 and ABC waveforms exhibited by *M. persicae*, a generalist aphid, feeding on the healthy *N. benthamiana* was 2.18 and 1.98 before and after fully developed infection symptoms, respectively. While, on *S. lycopersicum*, it was 1.24 and 0.67, respectively. This observation suggests that *N. benthamiana* is a more acceptable plant species for *M. persicae*. Furthermore, the observed trend shows an increased total time of the ABC waveform and a decrease in the total time of the E2 waveform on *S. lycopersicum*. This is due to the presence of CMV and satRNAs. In some cases, there is no engagement in the phloem phase. These factors suggest that aphids are more likely to abandon tomato plant. Consequently, they may transmit the virus and satRNA to neighbouring healthy plants (Carmo-Sousa et al., 2014). The change in aphid feeding behaviour caused by nc-satRNAs was observed in the case of *A. gossypii* during CMV acquisition on tomato plants, where aphids probed less willingly on CMV + nc-satRNA plants than in CMV-and CMV + non-nc-satRNA-inoculated plants (Escriu et al., 2000). Westwood et al. (2013) also found that CMV infection in *A. thaliana* induced an aphid feeding-deterrent 4-methoxy-indol-3-yl-methylglucosinolate, which inhibited phloem feeding. Moreover, previous studies have shown differences in the expression of genes associated with ethylene biosynthesis and its regulation, as well as alterations in the levels of proteins related to defense response to the insect in the virus and satRNA co-infected plants (Irian et al., 2007, Obrępalska-Stęplowska et al., 2015, Obrępalska-Stęplowska et al., 2018, Wrzesińska et al., 2018, Wrzesińska et al., 2021). Ethylene is one of the VOCs that was shown to play an important role in mediating the attraction of virus-infected plants to aphids (Bak et al., 2019, Mauck et al., 2014). It can therefore be hypothesised that satRNA presence may elicit stronger plants’ reaction to cucumoviral infection caused by the changes in the levels of the secondary metabolites, however this aspect should be explored in more detail.

In summary, our data provide for the first time a collective analysis of the influence of satRNA of cucumoviruses on the orientation and feeding behaviour of *M. persicae*, which may consequently result in differential virus acquisition and transmission efficiency. There are differences in the feeding behaviour caused by the presence of satRNAs, which may influence this efficiency. However, for feeding to occur, the aphid must be encouraged to move onto the infected plant, and this phase is crucial for the final transmission outcome. Here, we have shown that satRNAs that strongly exacerbated viral disease symptoms, by the example of PSV-G + satRNA and CMV + nc-satRNA, contributed to the reduced olfactory and feeding attractiveness of the host plants to *M. persicae*, which may explain, to some extent, the rarer occurrence of such satRNAs in the environment compared to those that attenuate symptoms. On the other hand, the satRNA genotypes that alleviate disease symptoms and reduce helper virus accumulation level, as in the case of PSV-P + satRNA, are beneficial for both virus and the host plant as the plants become more attractive to the insect vectors. This conclusion is in accordance with the statement that satRNA are considered as kind of buffers (Wrzesińska Krupa & Obrępalska Stęplowska, 2024). Under specific conditions, satRNAs are thought to stabilise the pathogenesis, which provide the survival of virus and host plant in the ecosystem. Whereas, in the instances where satRNA leads to symptom deterioration, it reduces the probability of virus transmission to the healthy plants.

## ACKNOWLEDGEMENTS

The authors would like to thank Prof. Xiaorong Tao and Prof. Fernando García-Arenal for kindly providing us with CMV-Fny agroinfectious clones and CMV satRNAs cDNA clones, respectively. This study was supported by National Science Centre (Poland) Grant no. 2018/29/N/NZ9/02467.

## STATEMENTS AND DECLARATIONS

The authors declare no conflict of interest. Data available on request from the authors.

---

**Figure S1.** Schematic representation of the olfactometer

**Table S1.** Primers used in this study for the synthesis of agroinfectious clones

**Table S2.** Primers used in this study for the detection of fragments corresponding to coat proteins (CPs) genes and the fragments of satRNAs (satRNAs) sequences of peanut stunt virus (PSV) and cucumber mosaic virus (CMV) using reverse transcription polymerase chain reaction (RT-PCR)

**Table S3.** Primers used in this study for the quantification of peanut stunt virus (PSV) and cucumber mosaic virus (CMV) genomic strands, and their satRNAs

**Table S4.** Statistical differences (treatment 1 vs treatment 2) between peanut stunt virus (PSV) genomic strands and satellite RNA (satRNA) accumulation level in *Nicotiana benthamiana* plants infected with PSV-G, PSV-G + satRNA, PSV-P, and PSV-P + satRNA according to Mann-Whitney *U* test

**Table S5.** Statistical differences (treatment 1 vs treatment 2) between cucumber mosaic virus (CMV) genomic strands and satellite RNA (satRNAs) accumulation level in *Nicotiana benthamiana* and *Solanum lycopersicum* plants infected with CMV with or without non-necrogenic satRNA (non-nc-satRNA) or necrogenic satRNA (nc-satRNA) according to Mann-Whitney *U* test

**Table S6.** Statistical differences between comparisons (treatment 1 vs treatment 2) of total duration (mean ± SEM) of electrical penetration graph (EPG) waveforms during *Myzus persicae* feeding on *Nicotiana benthamiana* plants infected with peanut stunt virus (PSV) strains (strain G or strain P) with or without satRNA before and after symptom development

**Table S7.** Statistical differences between comparisons (treatment 1 vs treatment 2) of total duration (mean ± SEM) of electrical penetration graph (EPG) waveforms during *Myzus persicae* feeding on *Nicotiana benthamiana* plants infected with cucumber mosaic virus (CMV) with or without non-necrogenic satRNA (non-nc-satRNA) or necrogenic satRNA (nc-satRNA) before and after symptom development

**Table S8.** Statistical differences between comparisons (treatment 1 vs treatment 2) of total duration (mean ± SEM) of electrical penetration graph (EPG) waveforms during *Myzus persicae* feeding on *Solanum lycopersicum* plants infected with cucumber mosaic virus (CMV) with or without non-necrogenic satRNA (non-nc-satRNA) or necrogenic satRNA (nc-satRNA) before and after symptom development

