## Supporting Information for "Symptom-modulating satRNAs of cucumoviruses affect the orientation and feeding behaviour of *Myzus persicae*"

Barbara Wrzesińska-Krupa^1^, Przemysław Strażyński^2^, Patryk Frąckowiak^1^, Aleksandra Obrępalska-Stęplowska^1^

^1^ Institute of Plant Protection – National Research Institute, Department of Molecular Biology and Biotechnology, Władysława Węgorka 20, 60-318 Poznań, Poland

^2^ Institute of Plant Protection – National Research Institute, Department of Entomology and Agricultural Pests, Władysława Węgorka 20, 60-318 Poznań, Poland

**Supporting Information**

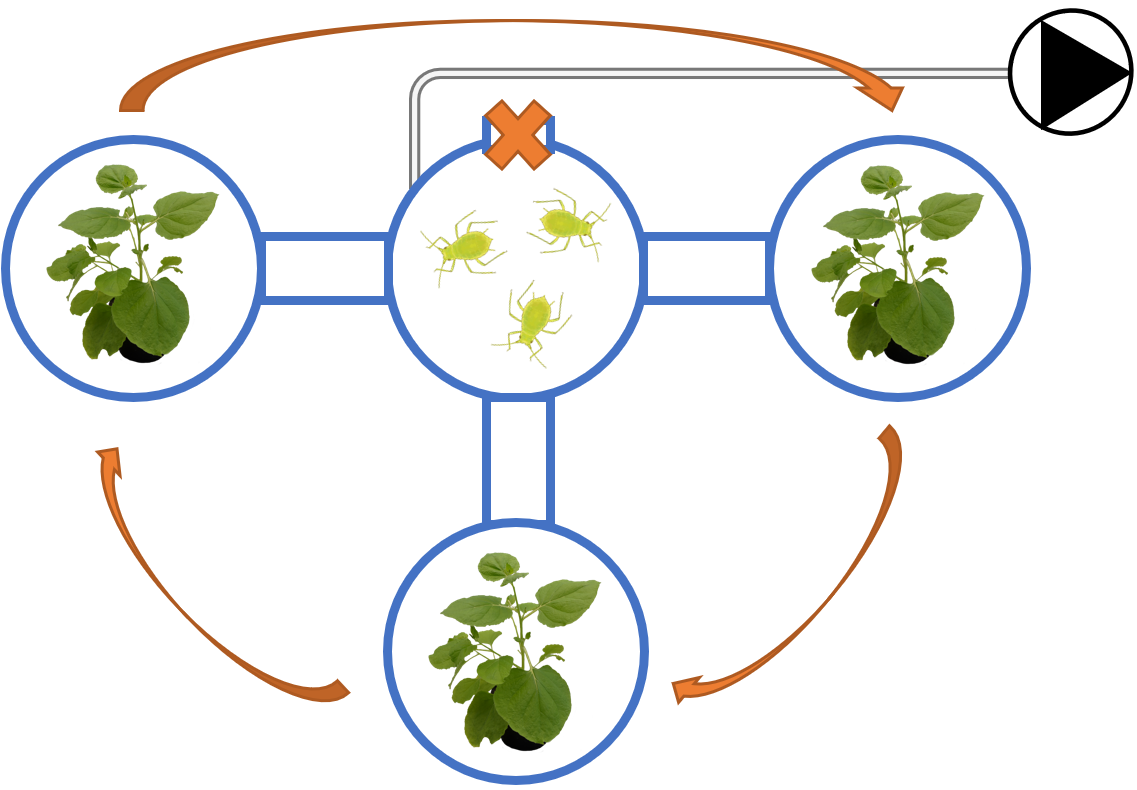

**Figure S1.** Schematic representation of the olfactometer. The arrows indicate the direction of the rotation of side jars. The
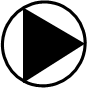
 sign stands for the air pump. Aphid image was obtained from TogoTV

**Table S1.** Primers used in this study for the synthesis of agroinfectious clones

| **Primer name** | **Primer sequence (5’→3’)** | **Function** |
| --- | --- | --- |
| PSVg1F | ATTTCATTTGGAGAGGGTTTTATCACGAGCGTACGG | Forward primer, for amplification of PSV-G RNA1; sequence essential for recombination with 5´ end of binary vector is underlined |
| PSVp2F | ATTTCATTTGGAGAGGGTTTTATCAAGAGCGTACGG | Forward primer, for amplification of PSV-G RNA2; sequence essential for recombination with 5´ end of binary vector is underlined (Wrzesińska et al., 2016) |
| PSVp3F | ATTTCATTTGGAGAGGGTTTTACCAACCAGGAATCTG | Forward primer, for amplification of PSV-G RNA3; sequence essential for recombination with 5´ end of binary vector is underlined (Wrzesińska et al., 2016) |
| PSVp123R | GATGCCATGCCGACCCTGGTCTCCTTATGGAACCCTC | Common reverse primer, for cDNA synthesis and amplification of PSV-G RNA1 and RNA3; sequence essential for recombination with 3´ end of binary vector is underlined (Wrzesińska et al., 2016) |
| PSVg2R | GATGCCATGCCGACCCTGGTCTCCTATGGAAACCCTC | Reverse primer, for cDNA synthesis and amplification of PSV-G RNA2; sequence essential for recombination with 3´ end of binary vector is underlined |
| CMVnonNCsatF | ATTTCATTTGGAGAGGGTTTTGTTTGTTAGAGAATT | Forward primer, for amplification of CMV non-necrogenic satRNA; sequence essential for recombination with 5´ end of binary vector is underlined |
| CMVnonNCsatR | GATGCCATGCCGACCCGGGTCCTGTAGAGGAATGTGTA | Reverse primer, for amplification of CMV non-necrogenic satRNA; sequence essential for recombination with 3´ end of binary vector is underlined |
| CMVNCsatF | ATTTCATTTGGAGAGGGTTTTGTTTGATGGAGAATT | Forward primer, for amplification of CMV necrogenic satRNA; sequence essential for recombination with 5´ end of binary vector is underlined |
| CMVNCsatR | GATGCCATGCCGACCCGGGTCCTGTAGAGGAATGATAG | Reverse primer, for amplification of CMV necrogenic satRNA; sequence essential for recombination with 3´ end of binary vector is underlined |

**Table S2.** Primers used in this study for the detection of fragments corresponding to coat proteins (CPs) genes and the fragments of satRNAs (satRNAs) sequences of peanut stunt virus (PSV) and cucumber mosaic virus (CMV) using reverse transcription polymerase chain reaction (RT-PCR)

| **Primer name** | **Primer sequence (5’→3’)** | **Annealing temperature** | **Function** |
| --- | --- | --- | --- |
| PSVCP | 1: TACCTTTTGGGTTCAATTCC | 45°C | Detection of PSV coat protein sequence (Obrępalska-Stęplowska et al., 2008) |
|  | 2: GACTGACCATTTTAGCCG |  |  |
| PSVsat | 1: CACTGTAATGGTGATGCGACAG | 55°C | Detection of PSV satRNA sequence |
|  | 2: CGGTACAGCAGTCCTGATAG |  |  |
| CMVcp | F: ATGGACAAATCTGAATCAACCAGT | 60°C | Detection of CMV coat protein sequence |
|  | R: TCAGACTGGGAGCACTCCA |  |  |
| CMVsat | F: GTTTTGTTTGATGGAGAATT | 52.7°C | Detection of CMV non-nc- and nc-satRNAs sequences |
|  | R: GGGTCCTGTAGAGGAATGATAG |  |  |

**Table S3.** Primers used in this study for the quantification of peanut stunt virus (PSV) and cucumber mosaic virus (CMV) genomic strands, and their satRNAs

| **Primer name** | **Primer sequence (5’→3’)** | **Annealing temperature** | **Function** |
| --- | --- | --- | --- |
| PSVq1 | F: CTTCTGCCCTCGTTGATAAAG | 57°C | Detection of PSV RNA1 in real-time PCR (Obrępalska-Stęplowska et al., 2015) |
|  | R: CATACCGATTTCGAATCACTT |  |  |
| PSVq2a | F: CTTCTAGGTATCCCCGTAAG | 60°C | Detection of PSV RNA2 in real-time PCR (Obrępalska-Stęplowska et al., 2015) |
|  | R: CAAGCACATTGATACCCTATC |  |  |
| PSVq3a | F: CTAGTCGGACTTTAACACAAC | 56°C | Detection of PSV RNA3 in real-time PCR (Obrępalska-Stęplowska et al., 2015) |
|  | R: ACGCTCATATATCCCTTAGAC |  |  |
| PARNA | 1: GGGAGGGCGGGCGTTCGTAGTG | 60°C | Detection of PSV satRNA in real-time PCR (Obrępalska-Stęplowska et al., 2015) |
|  | 2: GCCGTGGCCTTTCGTGGTC |  |  |
| CMVq1 | F: GGAGAGGAATGGGACGTGATATC | 60°C | Detection of CMV RNA1 in real-time PCR |
|  | R: CAAAATCTTCCCATCGGTAACAG |  |  |
| CMVq2a | F: ATGAGCTCCTTGTCGCTTTTG | 60°C | Detection of CMV RNA2 in real-time PCR |
|  | R: TTATTAAACGCAGGGCACCAT |  |  |
| CMVq3a | 1: CACGGTCGTATTGCTTCCTT | 60°C | Detection of CMV RNA2 in real-time PCR |
|  | 2: GAAACGCATTGCCCATCTAT |  |  |
| CMVqsat | F: GCGAGCTATGTCCGCTACTC | 60°C | Detection of CMV non-nc- and nc-satRNAs in real-time PCR |
|  | R: GACATTCACGGAGATCAGCA |  |  |

**Table S4.** Statistical differences (treatment 1 vs treatment 2) between peanut stunt virus (PSV) genomic strands and satellite RNA (satRNA) accumulation level in *Nicotiana benthamiana* plants infected with PSV-G, PSV-G + satRNA, PSV-P, and PSV-P + satRNA according to Mann-Whitney *U* test. * – *p*-value < 0.05

| **Genomic strand** | **PSV** | | |
| --- | --- | --- | --- |
|  | **Treatment 1** | **Treatment 2** | ***p*-value** |
| RNA1 | PSV-G | PSV-G + satRNA | 0.724 |
|  | PSV-P | PSV-P + satRNA | 0.0217* |
| RNA2 | PSV-G | PSV-G + satRNA | 0.157 |
|  | PSV-P | PSV-P + satRNA | 0.000407* |
| RNA3 | PSV-G | PSV-G + satRNA | 0.724 |
|  | PSV-P | PSV-P + satRNA | 0.000404* |
| satRNA | PSV-G + satRNA | PSV-P + satRNA | 0.25 |

**Table S5.** Statistical differences (treatment 1 vs treatment 2) between cucumber mosaic virus (CMV) genomic strands and satellite RNA (satRNAs) accumulation level in *Nicotiana benthamiana* and *Solanum lycopersicum* plants infected with CMV with or without non-necrogenic satRNA (non-nc-satRNA) or necrogenic satRNA (nc-satRNA) according to Mann-Whitney *U* test. * – *p*-value < 0.05

| **Genomic strand** | **CMV – *N. benthamiana*** | | | **CMV – *S. lycopersicum*** | | |
| --- | --- | --- | --- | --- | --- | --- |
|  | **Treatment 1** | **Treatment 2** | ***p*-value** | **Treatment 1** | **Treatment 2** | ***p*-value** |
| RNA1 | CMV | CMV + non-nc-satRNA | 0.000412* | CMV | CMV + non-nc-satRNA | 0.000409* |
|  | CMV | CMV + nc-satRNA | 0.000409* | CMV | CMV + nc-satRNA | 0.000412* |
|  | CMV + non-nc-satRNA | CMV + nc-satRNA | 0.0009409* | CMV + non-nc-satRNA | CMV + nc-satRNA | 0.596 |
| RNA2 | CMV | CMV + non-nc-satRNA | 0.000404* | CMV | CMV + non-nc-satRNA | 0.000409* |
|  | CMV | CMV + nc-satRNA | 0.000404* | CMV | CMV + nc-satRNA | 0.000409* |
|  | CMV + non-nc-satRNA | CMV + nc-satRNA | 0.000412* | CMV + non-nc-satRNA | CMV + nc-satRNA | 0.2 |
| RNA3 | CMV | CMV + non-nc-satRNA | 0.251 | CMV | CMV + non-nc-satRNA | 0.000409* |
|  | CMV | CMV + nc-satRNA | 0.000398* | CMV | CMV + nc-satRNA | 0.000409* |
|  | CMV + non-nc-satRNA | CMV + nc-satRNA | 0.000401* | CMV + non-nc-satRNA | CMV + nc-satRNA | 0.757 |
| satRNA | CMV + non-nc-satRNA | CMV + nc-satRNA | 0.000412* | CMV + non-nc-satRNA | CMV + nc-satRNA | 0.00611* |

**Table S6.** Statistical differences between comparisons (treatment 1 vs treatment 2) of total duration (mean ± SEM) of electrical penetration graph (EPG) waveforms during *Myzus persicae* feeding on *Nicotiana benthamiana* plants infected with peanut stunt virus (PSV) strains (strain G or strain P) with or without satRNA before and after symptom development. Np - non-penetration, ABC – penetration of peripheral tissues (mesophyll), E1 – pricking phloem and secretion of saliva, E2 – collection of phloem sap. Significant differences were calculated according to Mann-Whitney *U* test. * – *p*-value < 0.05

| **EPG waveform** | **PSV strain G** | | | **PSV strain P** | | |
| --- | --- | --- | --- | --- | --- | --- |
|  | **Treatment 1** | **Treatment 2** | ***p*-value** | **Treatment 1** | **Treatment 2** | ***p*-value** |
| Before symptom development | | | | | | |
| Np | MOCK | PSV-G | 0.637 | MOCK | PSV-P | 0.713 |
|  | MOCK | PSV-G + satRNA | 0.0086* | MOCK | PSV-P + satRNA | 0.156 |
|  | PSV-G | PSV-G + satRNA | 0.372 | PSV-P | PSV-P + satRNA | 0.27 |
| ABC | MOCK | PSV-G | 0.0313* | MOCK | PSV-P | 0.27 |
|  | MOCK | PSV-G + satRNA | 0.875 | MOCK | PSV-P + satRNA | 0.0101* |
|  | PSV-G | PSV-G + satRNA | 0.0239* | PSV-P | PSV-P + satRNA | 0.0239* |
| E1 | MOCK | PSV-G | 0.636 | MOCK | PSV-P | 0.495 |
|  | MOCK | PSV-G + satRNA | 0.27 | MOCK | PSV-P + satRNA | 0.227 |
|  | PSV-G | PSV-G + satRNA | 0.713 | PSV-P | PSV-P + satRNA | 0.834 |
| E2 | MOCK | PSV-G | 0.958 | MOCK | PSV-P | 1 |
|  | MOCK | PSV-G + satRNA | 0.564 | MOCK | PSV-P + satRNA | 0.528 |
|  | PSV-G | PSV-G + satRNA | 0.713 | PSV-P | PSV-P + satRNA | 0.713 |
| After symptom development | | | | | | |
| Np | MOCK | PSV-G | 0.0406* | MOCK | PSV-P | 0.0181* |
|  | MOCK | PSV-G + satRNA | 0.000939* | MOCK | PSV-P + satRNA | 0.0000939* |
|  | PSV-G | PSV-G + satRNA | 0.431 | PSV-P | PSV-P + satRNA | 0.0313* |
| ABC | MOCK | PSV-G | 0.00136* | MOCK | PSV-P | 0.00195* |
|  | MOCK | PSV-G + satRNA | 0.87 | MOCK | PSV-P + satRNA | 0.000939* |
|  | PSV-G | PSV-G + satRNA | 0.0661 | PSV-P | PSV-P + satRNA | 0.156 |
| E1 | MOCK | PSV-G | 0.00388* | MOCK | PSV-P | 0.0181* |
|  | MOCK | PSV-G + satRNA | 0.0023* | MOCK | PSV-P + satRNA | 0.00193* |
|  | PSV-G | PSV-G + satRNA | 0.713 | PSV-P | PSV-P + satRNA | 0.4 |
| E2 | MOCK | PSV-G | 0.104 | MOCK | PSV-P | 0.318 |
|  | MOCK | PSV-G + satRNA | 0.0406* | MOCK | PSV-P + satRNA | 0. 313* |
|  | PSV-G | PSV-G + satRNA | 0.958 | PSV-P | PSV-P + satRNA | 0.0661 |

**Table S7.** Statistical differences between comparisons (treatment 1 vs treatment 2) of total duration (mean ± SEM) of electrical penetration graph (EPG) waveforms during *Myzus persicae* feeding on *Nicotiana benthamiana* plants infected with cucumber mosaic virus (CMV) with or without non-necrogenic satRNA (non-nc-satRNA) or necrogenic satRNA (nc-satRNA) before and after symptom development. Np - non-penetration, ABC – penetration of peripheral tissues (mesophyll), E1 – pricking phloem and secretion of saliva, E2 – collection of phloem sap. Significant differences were calculated according to Mann-Whitney *U* test. * – *p*-value < 0.05

| **EPG waveform** | **Treatment 1** | **Treatment 2** | ***p*-value** | **EPG waveform** | **Treatment 1** | **Treatment 2** | ***p*-value** |
| --- | --- | --- | --- | --- | --- | --- | --- |
| Before symptom development | | | | After symptom development | | | |
| Np | MOCK | CMV | 0.793 | Np | MOCK | CMV | 0.27 |
|  | MOCK | CMV + non-nc-satRNA | 0.318 |  | MOCK | CMV + non-nc-satRNA | 0.0831 |
|  | MOCK | CMV + nc-satRNA | 0.372 |  | MOCK | CMV + nc-satRNA | 0.0239* |
|  | CMV | CMV + non-nc-satRNA | 0.372 |  | CMV | CMV + non-nc-satRNA | 0.495 |
|  | CMV | CMV + nc-satRNA | 0.4 |  | CMV | CMV + nc-satRNA | 0.372 |
| ABC | MOCK | CMV | 1 | ABC | MOCK | CMV | 0.372 |
|  | MOCK | CMV + non-nc-satRNA | 0.564 |  | MOCK | CMV + non-nc-satRNA | 0.564 |
|  | MOCK | CMV + nc-satRNA | 0.318 |  | MOCK | CMV + nc-satRNA | 0.495 |
|  | CMV | CMV + non-nc-satRNA | 0.495 |  | CMV | CMV + non-nc-satRNA | 0.713 |
|  | CMV | CMV + nc-satRNA | 0.156 |  | CMV | CMV + nc-satRNA | 0.637 |
| E1 | MOCK | CMV | 0.875 | E1 | MOCK | CMV | 0.793 |
|  | MOCK | CMV + non-nc-satRNA | 1 |  | MOCK | CMV + non-nc-satRNA | 0.188 |
|  | MOCK | CMV + nc-satRNA | 0.462 |  | MOCK | CMV + nc-satRNA | 0.0356* |
|  | CMV | CMV + non-nc-satRNA | 0.637 |  | CMV | CMV + non-nc-satRNA | 0.371 |
|  | CMV | CMV + nc-satRNA | 0.431 |  | CMV | CMV + nc-satRNA | 0.127 |
| E2 | MOCK | CMV | 0.637 | E2 | MOCK | CMV | 0.793 |
|  | MOCK | CMV + non-nc-satRNA | 0.431 |  | MOCK | CMV + non-nc-satRNA | 0.564 |
|  | MOCK | CMV + nc-satRNA | 0.27 |  | MOCK | CMV + nc-satRNA | 0.227 |
|  | CMV | CMV + non-nc-satRNA | 0.637 |  | CMV | CMV + non-nc-satRNA | 0.713 |
|  | CMV | CMV + nc-satRNA | 0.27 |  | CMV | CMV + nc-satRNA | 0.227 |

**Table S8.** Statistical differences between comparisons (treatment 1 vs treatment 2) of total duration (mean ± SEM) of electrical penetration graph (EPG) waveforms during *Myzus persicae* feeding on *Solanum lycopersicum* plants infected with cucumber mosaic virus (CMV) with or without non-necrogenic satRNA (non-nc-satRNA) or necrogenic satRNA (nc-satRNA) before and after symptom development. Np - non-penetration, ABC – penetration of peripheral tissues (mesophyll), E1 – pricking phloem and secretion of saliva, E2 – collection of phloem sap. Significant differences were calculated according to Mann-Whitney *U* test. * – *p*-value < 0.05

| **EPG waveform** | **Treatment 1** | **Treatment 2** | ***p*-value** | **EPG waveform** | **Treatment 1** | **Treatment 2** | ***p*-value** |
| --- | --- | --- | --- | --- | --- | --- | --- |
| Before symptom development | | | | After symptom development | | | |
| Np | MOCK | CMV | 0.958 | Np | MOCK | CMV | 0.637 |
|  | MOCK | CMV + non-nc-satRNA | 0.318 |  | MOCK | CMV + non-nc-satRNA | 0.156 |
|  | MOCK | CMV + nc-satRNA | 0.958 |  | MOCK | CMV + nc-satRNA | 0.248 |
|  | CMV | CMV + non-nc-satRNA | 0.431 |  | CMV | CMV + non-nc-satRNA | 0.431 |
|  | CMV | CMV + nc-satRNA | 0.958 |  | CMV | CMV + nc-satRNA | 0.495 |
| ABC | MOCK | CMV | 0.564 | ABC | MOCK | CMV | 0.564 |
|  | MOCK | CMV + non-nc-satRNA | 0.495 |  | MOCK | CMV + non-nc-satRNA | 0.293 |
|  | MOCK | CMV + nc-satRNA | 0.495 |  | MOCK | CMV + nc-satRNA | 0.27 |
|  | CMV | CMV + non-nc-satRNA | 0.227 |  | CMV | CMV + non-nc-satRNA | 0.495 |
|  | CMV | CMV + nc-satRNA | 0.27 |  | CMV | CMV + nc-satRNA | 0.958 |
| E1 | MOCK | CMV | 0.793 | E1 | MOCK | CMV | 1 |
|  | MOCK | CMV + non-nc-satRNA | 0.189 |  | MOCK | CMV + non-nc-satRNA | 0.225 |
|  | MOCK | CMV + nc-satRNA | 0.248 |  | MOCK | CMV + nc-satRNA | 0.752 |
|  | CMV | CMV + non-nc-satRNA | 0.495 |  | CMV | CMV + non-nc-satRNA | 0.292 |
|  | CMV | CMV + nc-satRNA | 0.495 |  | CMV | CMV + nc-satRNA | 0.431 |
| E2 | MOCK | CMV | 0.563 | E2 | MOCK | CMV | 0.528 |
|  | MOCK | CMV + non-nc-satRNA | 0.189 |  | MOCK | CMV + non-nc-satRNA | 0.263 |
|  | MOCK | CMV + nc-satRNA | 0.103 |  | MOCK | CMV + nc-satRNA | 0.161 |
|  | CMV | CMV + non-nc-satRNA | 0.431 |  | CMV | CMV + non-nc-satRNA | 0.122 |
|  | CMV | CMV + nc-satRNA | 0.834 |  | CMV | CMV + nc-satRNA | 0.0845 |
